# Auranofin Inhibition of Thioredoxin Reductase Sensitizes Lung Neuroendocrine Tumor Cells (NETs) and Small Cell Lung Cancer (SCLC) Cells to Sorafenib as well as Inhibiting SCLC Xenograft Growth

**DOI:** 10.1101/2023.05.07.539772

**Authors:** Spenser S. Johnson, Dijie Liu, Jordan T. Ewald, Claudia Robles-Planells, Keegan A. Christensen, Khaliunaa Bayanbold, Brian R. Wels, Shane R. Solst, M. Sue O’Dorisio, Bryan G. Allen, Yusuf Menda, Douglas R. Spitz, Melissa A. Fath

## Abstract

Thioredoxin Reductase (TrxR) is a key enzyme in hydroperoxide detoxification through peroxiredoxin enzymes and in thiol-mediated redox regulation of cell signaling. Because cancer cells produce increased steady-state levels of reactive oxygen species (ROS; i.e., superoxide and hydrogen peroxide), TrxR is currently being targeted in clinical trials using the anti-rheumatic drug, auranofin (AF). AF treatment decreased TrxR activity and clonogenic survival in small cell lung cancer (SCLC) cell lines (DMS273 and DMS53) as well as the lung atypical (neuroendocrine tumor) NET cell line H727. AF treatment also significantly sensitized DMS273 and H727 cell lines *in vitro* to sorafenib, a multi-kinase inhibitor that was shown to decrease intracellular glutathione. The pharmacokinetic and pharmacodynamic properties of AF treatment in a mouse SCLC xenograft model was examined to maximize inhibition of TrxR activity without causing toxicity. AF was administered intraperitoneally at 2 mg/kg or 4 mg/kg (IP) once (QD) or twice daily (BID) for 1 to 5 days in mice with DMS273 xenografts. Plasma levels of AF were 10-20 μM (determined by mass spectrometry of gold) and the optimal inhibition of TrxR (50 %) was obtained at 4 mg/kg once daily, with no effect on glutathione peroxidase 1 activity. When this daily AF treatment was extended for 14 days a significant prolongation of median survival from 19 to 23 days (p=0.04, N=30 controls, 28 AF) was observed without causing changes in animal bodyweight, CBCs, bone marrow toxicity, blood urea nitrogen, or creatinine. These results show that AF is an effective inhibitor of TrxR both *in vitro* and *in vivo* in SCLC, capable of sensitizing NETs and SCLC to sorafenib, and supports the hypothesis that AF could be used as an adjuvant therapy with agents known to induce disruptions in thiol metabolism to enhance therapeutic efficacy.

## Introduction

Pulmonary neuroendocrine tumors and carcinomas are a group of heterogeneous neoplasms that are classified according to their degree of differentiation, mitosis, and degree of necrosis.^1^ For lung neuroendocrine tumors (NET), the 5-year survival is 35% for patients presenting with distant metastases, which is the initial presentation in 35% of patients. The survival rate drops to 4% in poorly differentiated NETs.^2^ The average survival time for patients with lung neuroendocrine carcinomas (NEC) (including small cell lung cancer [SCLC]) is 10 months, with a 2-year survival of only 6%. ^3,4^ Therefore, improved treatment options are of utmost importance for pulmonary NETs and NECs.

A hallmark of cancer is the increased steady-state levels of reactive oxygen species (ROS) often stemming from one electron reductions of O_2_ in mitochondrial electron transport chains to form superoxide radical (O_2_^·-^) which is rapidly converted to H_2_O_2_^5^ Cancer cells adapt to increased levels of ROS by upregulating endogenous antioxidant systems, including the scavenging enzymes superoxide dismutase and catalase, as well as enzymes in the glutathione-dependent and thioredoxin-dependent hydroperoxide metabolic pathways. The glutathione and thioredoxin systems play key roles in the overall cellular oxidation state because of their intracellular dithiol recycling reactions in hydroperoxide metabolism and redox signaling. ^6^ In particular, the thioredoxin system consists of the redox-active proteins thioredoxin (Trx) and the NADPH electron acceptor thioredoxin reductase (TrxR) which primarily function to reduce peroxiredoxin (Prx) during the catalytic reduction of hydroperoxides. TrxR has three known mammalian isoforms that can be found in the cytosol, mitochondria, and one specific to spermatozoa.^7^ According to the Project Score database, knock-out of TrxR has shown an anti-tumoral effect in 21% of 324 cell-lines subjected to CRISPR-Cas9 screens ^8^ clearly indicating TrxR as a potential target for cancer treatment. In addition, Bebber *et al*. found that overall survival of SCLC patients with low TrxR expression was significantly better than patients with tumors with high TrxR expression.^9^ These data provide the rationale for targeting TrxR in SCLC.

Auranofin (AF), a gold phosphine salt, was developed as a rheumatic drug as it was originally thought to exert its effects by inhibiting humoral immunity. The most common acute adverse reactions associated with AF treatment were gastrointestinal with loose stools or diarrhea occurring in 39% of patients. Rashes and pruritis were also common, occurring in >10% of patients.^10-12^ In addition, chronic use of gold salts caused thrombocytopenia in 0.7% of patients.^13^ Although diarrhea and stomach upset occurred in the initial month of treatment, other adverse events typically occurred only after prolonged therapy.^14^ Clinical AF use has declined due to the availability of more effective and well-tolerated biologic treatments for rheumatoid arthritis.

However, AF was also discovered to irreversibly inhibit both cytosolic and mitochondrial TrxR.^15,16^ This discovery has led to the active investigation of repurposing AF as an anticancer agent. Currently, four clinical trials using AF are underway for chronic lymphocytic leukemia (NCT01419691, NCT01747798), ovarian cancer (NCT03456700), and recurrent lung cancer (NCT01737502). We have previously shown simultaneous disruption of glutathione and thioredoxin metabolism enhanced non-small cell lung cancer cell killing and sensitivity to chemotherapy.^17^ In addition, we demonstrated that treatment of breast cancer cells with AF before external beam radiation, decreased the number of invading metastatic breast cancer cells.^18^ We also demonstrated that suppressing TrxR with AF can sensitize breast cancer stem cells to ROS induced stem cell transitions associated with EMT and cytotoxicity associated with 2-deoxyglucose treatment.^19^

The current study examined the effects of AF in lung NETs and SCLCs *in vitro* as well as in responses to sorafenib, a multi-kinase inhibitor that is known to decrease intracellular glutathione and sensitize non-small cell lung cancer to cisplatin.^20^ In addition, because very little is known about the pharmacokinetics of AF in mice, we developed a SCLC xenograft model and determined a safe and efficacious dose of AF that significantly inhibited TrxR in tumors without affecting glutathione peroxidase 1. After a two-week dosing schedule of AF at 4 mg/kg IP once daily, complete blood counts as well as hematopoietic stem cell and progenitor cell populations were measured to show AF was well-tolerated by the hematopoietic system of tumor bearing mice. Renal and liver function, evaluated by serum chemistry analyses, as well as weight loss and animal body conditioning also indicated AF was well-tolerated in this model system. Finally, median overall survival of the mice with SCLC was significantly prolonged by treatment with AF. These results provide a rigorous characterization of the effects of AF on a SCLC xenograft mouse model and continue to support the hypothesis that repurposing of AF as an adjuvant to cancer therapy in combination with agents that disrupt thiol metabolism is justified.

## Methods and Materials

### Cell Culture

DMS53 human small cell lung cancer cells (Aka. Neuroendocrine Carcinoma Cells) (NECs) and H727 human (Aka. lung neuroendocrine tumor cells) (NETs) were obtained from American Type Culture Collection (ATCC). DMS273 human small cell lung cancer cells (NECs) were obtained from the European Collection of Authenticated Cell Cultures were obtained from Millipore-Sigma. Cells were maintained in RPMI 1640 media (Mediatech Inc., Manassas, VA) with 10% fetal bovine serum (Atlanta Biologicals, Lawrenceville, GA). Cultures were maintained in 5% CO_2_ and humidified in a 37°C incubator. Cells were used within 20 passages from ATCC and tested as mycoplasma negative.

### Clonogenic Cell Survival Assay

Exponentially growing DMS273, DMS53 and H727 cells on 60 mm dishes were treated with increasing doses of sorafenib (Cayman Chemicals) for 48 hours with or without 1 μM AF for the last 3 hours. Floating cells in medium were collected and combined with the attached cells from the same dish that were trypsinized with 1 ml trypsin-EDTA (CellGro, Herndon, VA). Samples were centrifuged and cells were counted using a Beckman Coulter Counter. Cells were plated at low density (150-20,000 cells per dish), and clones were allowed to grow for 10-14 days in maintenance media with 0.1% gentamycin added. Cells were then fixed with 70% ethanol and stained with Coomassie blue for analysis of clonogenic survival. Individual assay colony counts were normalized to that of control with at least 3 cloning dishes per condition, repeated in at least 3 separate experiments.

### Thioredoxin Reductase Assay

TrxR activity was determined spectrophotometrically using the reduction of 5,5’-dithiobis(2-nitrobenzoic) acid (DTNB) with NADPH to 5-thio-2-nitrobenzoic acid (TNB), which generates a yellow color at 412 nm (Millipore-Sigma CS0170). Enzymatic activity was determined by subtracting the time dependent increase in absorbance at 412 nm in the presence of the TrxR activity inhibitor, aurothioglucose from total activity. One unit of activity was defined as 1 μM TNB formed/(min·mg protein). Protein concentrations were determined by the Lowry assay.

### Glutathione Assay

Exponentially growing DMS273 and H727 cells on 100 mm dishes were treated for 48 hours with sorafenib. Immediately following treatment, the cultures were washed in cold PBS then the cells were scrape harvested in 300 μL 5% 5-sulfosalicylic acid (Sigma) in water and stored at -20°C for a maximum of 72 hours. Total glutathione (GSH) content was determined by the method of Griffith.^21^ The rates of the reaction were compared to similarly prepared GSH standard curves. Glutathione determinations were normalized to the protein content of the insoluble pellet from the SSA extracts 2.5% SDS in 0.1 N bicarbonate using the BCA Protein Assay Kit (Thermo Scientific, Rockford, IL).

### Glutathione peroxidase 1 (GPx1) Activity Assay

Glutathione peroxidase was determined spectrophotometrically using the method of Lawrence and Burk as described previously.^22^ Enzymatic activity was determined by measuring the disappearance of NADPH at 340 nm in the presence of GPx1 from cell pellets or standards. One unit of activity was defined as 1 µmole NAPDH oxidized/min at specified glutathione (GSH) concentration. Protein concentrations were determined by the Lowry assay.

### Animal Experiments

Female 6–8-week-old athymic-nu/nu nude mice were purchased from Envigo. Female mice were used for these studies as they have been shown to be more sensitive to the toxic effects of AF.^23^ Mice were housed in a pathogen-free barrier room in the Animal Care Facility at the University of Iowa and handled using aseptic procedures. All procedures were approved by the IACUC committee of the University of Iowa and conformed to the guidelines established by NIH. Mice were allowed at least one week to acclimate prior to beginning experimentation; food and water were made freely available. Tumor cells were inoculated into nude mice by subcutaneous injection of 0.1 mL aliquots of saline containing 1 × 10^6^ DMS273 cells into the right and/or left flank using 25-gauge needles. Mouse health and activity levels were monitored daily during drug treatment as per our IACUC approved protocol. Mice were terminally anesthetized using 200 µl IP injection of ketamine/xylazine 17.5/2.5 µg/µl mix. When analgesia was achieved, as determined by lack of toe pinch response, blood was withdrawn via cardiac puncture followed by cervical dislocation as per our IACUC approved protocol.

### *In vivo* AF Dose Escalation

In the pilot dose escalation study to determine what AF dosing scheme gave the most rapid and durable inhibition of TrxR activity based on our previous work ^18,19^, five groups of six mice/group (DMS273 tumors growing on both flanks; 2 tumors/mouse) were given AF IP once (QD) or twice (BID) a day for 1 or 5 days. We sacrificed half of the mice (3 mice/group) after 24 hr to determine if the effects of AF on TrxR activity could be seen quickly (**Figure 2A-C**). The other 3 mice per group were sacrificed after 5 days as it has been demonstrated in humans that gold levels in plasma reach a 1 µM steady-state after several doses (a level of AF shown to inhibit TrxR *in vitro*).^24^ Group 1 was given vehicle injection; group 2 was given 2 mg/kg AF QD; group 3 was given 2 mg/kg AF BID; group 4 was given 4 mg/kg AF QD; and group 5 was given 4 mg/kg AF BID. AF was made by dissolving 5.0 mg AF in 250 µl of absolute ethanol in a 10 ml glass beaker. When completely dissolved 250 µl Kolliphor EL (Sigma C5135) and 4 ml of 0.9% normal saline for injection (Hospira) were added and pH was adjusted to 7-8 range using approximately 25 µl 8.4% sodium bicarbonate for injection (Hospira) followed by addition of normal saline for injection to adjust final volume to 8.33 ml and filtered using 200-micron filter into a sterile vial. The final AF concentration was 0.6 mg/ml and it was made fresh daily. Mice in the control group were administered vehicle without drug. Mice were euthanized 21 hours after the last QD dose and 9 hours after the last BID dose; tumors and organs were frozen in liquid nitrogen for later evaluation for TrxR activity. Terminal blood was collected via cardiac puncture under anesthesia and plasma separated via centrifugation at 10,000 rcf for 10 min.

### Determining Gold content in Plasma

Plasma samples collected 9 hours after final BID dose (5 days of therapy) and gold content determined were prepared by treatment with aqua regia followed by dilution and analysis using inductively coupled plasma mass spectrometry (ICPMS). Aqua regia was prepared from Optima grade (Fisher Scientific) hydrochloric and nitric acids (3:1, respectively). A 100-µL aliquot of plasma was transferred to a 15-mL centrifuge tube, 500 μL aqua regia added and the mixture heated on a hot block at 90 +/-4 °C for 30 minutes. After cooling, volume was brought to 2.0 mL using deionized water. A 500-μL aliquot of the digestate was mixed with an equal volume of internal standard (10 μg/L Iridium) and analyzed using ICPMS (Agilent 7700) with an external calibration curve.

### Tumor growth and survival

In the survival experiments, 3 cohorts of mice (randomly assigned groups of 9-10 mice/group bearing DMS273 flank tumors) were administered AF at 4 mg/kg or vehicle interperitoneally (IP) once a day for 14 consecutive days (**Figure 5 and supplemental Figure 1**). When xenograft tumor volumes measured approximately 100 mm^3^ treatments were started. Additionally, in the first cohort two mice in the AF group and two mice in the control group (each bearing two tumors), were euthanized 24 hours after the last dose of 4 mg/kg AF (day 15), and tumors were frozen at -80 °C to determine TrxR activity assay. Similarly, in the third experiment 4 mice in the control group and 4 mice in the AF group were likewise sacrificed on day 15 to determine TrxR activity (**Figure 3**). On day 15, blood was obtained via mandibular vein and diluted 1:10 in PBS and a complete blood cell count (CBC) processed in a Siemens ADVIA 120 Hematology Analyzer. The remaining mice were monitored for signs of toxicity by examining body weight and general activity level, and tumors measured daily by a blinded investigator using Vernier calipers (Vol = (Length x Width^2^)/2). Mice were euthanized on the day when tumor size exceeded 1000 mm^3^ for 2 consecutive days (a sized determined to not cause discomfort or mobility issues in the mice). No mice died before reaching this endpoint. Blood was collected from a select group of 7 control and 5 AF mice from the third cohort via cardiac puncture. Biochemical assays for AST, ALT, BUN, and creatinine were standard systemic toxicity biomarker assays performed commercially by Antech Diagnostics using protocols described on their website https://www.antechdiagnostics.com/test-guide/ (**Figure 4D-L**). <u>Bone marrow harvest and flow cytometry analysis</u> Immediately after euthanasia, whole bone marrow cells were obtained from 9 AF treated and 13 control mice per group, by flushing one femur with PBS 2% FBS followed by erythrocyte elimination with ACK Lysing Buffer (10-548E, BioWhittaker, Lonza) (**Figure 4BC**). The remaining cells were filtered through 40 μm pore size cell strainer and counted. 1x10^6^ cells were stained with Zombie Aqua (423102, BioLegend) for 20 min at room temperature for live/dead cells discrimination, followed by a 30 min surface staining on ice using PBS 1% BSA as staining buffer with allophycocyanin (APC)-conjugated, lineage negative (Lin-) cocktail (BD Bioscience), PerCP/Cy5.5-conjugated anti-Sca1 (Clone D7, BioLegend), APC/Cy7-conjugated anti-cKit (Clone 2B8, BioLegend), PE-conjugated anti-CD135 (Clone A2F10, BioLegend), FITC-conjugated anti-CD48 (Clone HM48-1, BioLegend), PE/Cy7-conjugated anti-CD150 (Clone TC15-12F12.2, BioLegend) and BV421-conjugated anti-CXCR4 (Clone L276F12, BioLegend). Cells were immediately analyzed using an LSR II flow cytometer (Becton Dickinson). The biomarkers were applied to identify multipotent hematopoietic stem cells (HSC) (Lin-cKit+ Sca1+, LSK), ST-HSC (short-term hematopoietic stem cells, Lin-cKit+ Sca1+ CD135-CD48-CD150-), LT-HSC (long-term hematopoietic stem cells, Lin-cKit+ Sca1+ CD135-CD48+CD150-) (ST-HSC/LT-HSC and committed progenitors (hematopoietic stem and progenitor cells, HSPC, Lin-cKit+ Sca1+ CD135-) using FlowJo Software (Treestar, Ashland, OR). <u>Statistical analysis</u> The biochemical data analysis was performed using ANOVA multiple comparisons with uncorrected Fishers LSD on Graph Pad Prism software. The *in vivo* analysis regression analysis was used to model tumor growth as a nonlinear function of follow-up time and to make treatment group comparisons of estimated tumor means. Survival graphs were obtained with the methods of Kaplan–Meier and compared with log-rank tests.

## Results

### AF inhibits TrxR and decreases clonogenic survival in lung NET and NECs via a thiol mediated mechanism

We and others have demonstrated that treatment with physiologically relevant doses of AF decreases TrxR activity and sensitizes a variety of cancer cells to chemotherapy and ionizing radiation both *in vitro* and *in vivo.* ^17,18,25,26^ Recently it was demonstrated that a SCLC subpopulation with neuroendocrine like phenotype was resistant to cell death from lipid oxidation and also demonstrated a clear dependence on the Trx driven anti-oxidant pathway for survival.^9^ We also have demonstrated that a subpopulation of breast cancer stem cells was more dependent of the Trx system.^19^ To extend these observations, we treated two SCLC cell lines (DMS53, DMS273) (NECs) and one lung NET cell line (H727) with AF for 3 h followed by the TrxR activity assay and clonogenic survival assay. **Figure 1AB** demonstrates that 1 μM AF for 24 hr significantly decreases TrxR activity and clonogenic survival in lung NET and NECs. We have previously demonstrated in lung, prostate, and breast cancer that simultaneous inhibition of Trx- and glutathione-dependent pathways of hydroperoxide metabolism resulted in enhanced cell death via a mechanism of thiol-mediated oxidative stress. ^17-19,27^ It has also been demonstrated in SCLC cells that targeting both the glutathione and Trx-dependent pathways simultaneously prevent cancer stem cell plasticity and improves overall survival.^9^ Others have found that the clinically approved anti-cancer drug sorafenib inhibits system Xct (-) function resulting in decreased intracellular glutathione and an increase in lipid oxidation stress.^28^

Therefore, we hypothesized that combining AF and sorafenib would decrease NEC and NET cell survival. Initially, we treated H727, and DMS273 cells with pharmacologically relevant doses of sorafenib followed by quantification of intercellular glutathione verifying a dose dependent reduction of total intracellular glutathione in each cell line (**Figure 1C**). To test combination of AF + sorafenib, all cell lines were grown in the exponential phase then given escalating doses of sorafenib for 48 hours with or without AF at the final 24 hours. AF significantly enhanced clonogenic cell death of sorafenib in both DMS273 and H727 cells (**Figure 1DE**). These results underline the importance of the hydroperoxide detoxification pathways in cancer cells as well as targeting both major thiol mediated-detoxification pathways.

**Figure 1:**
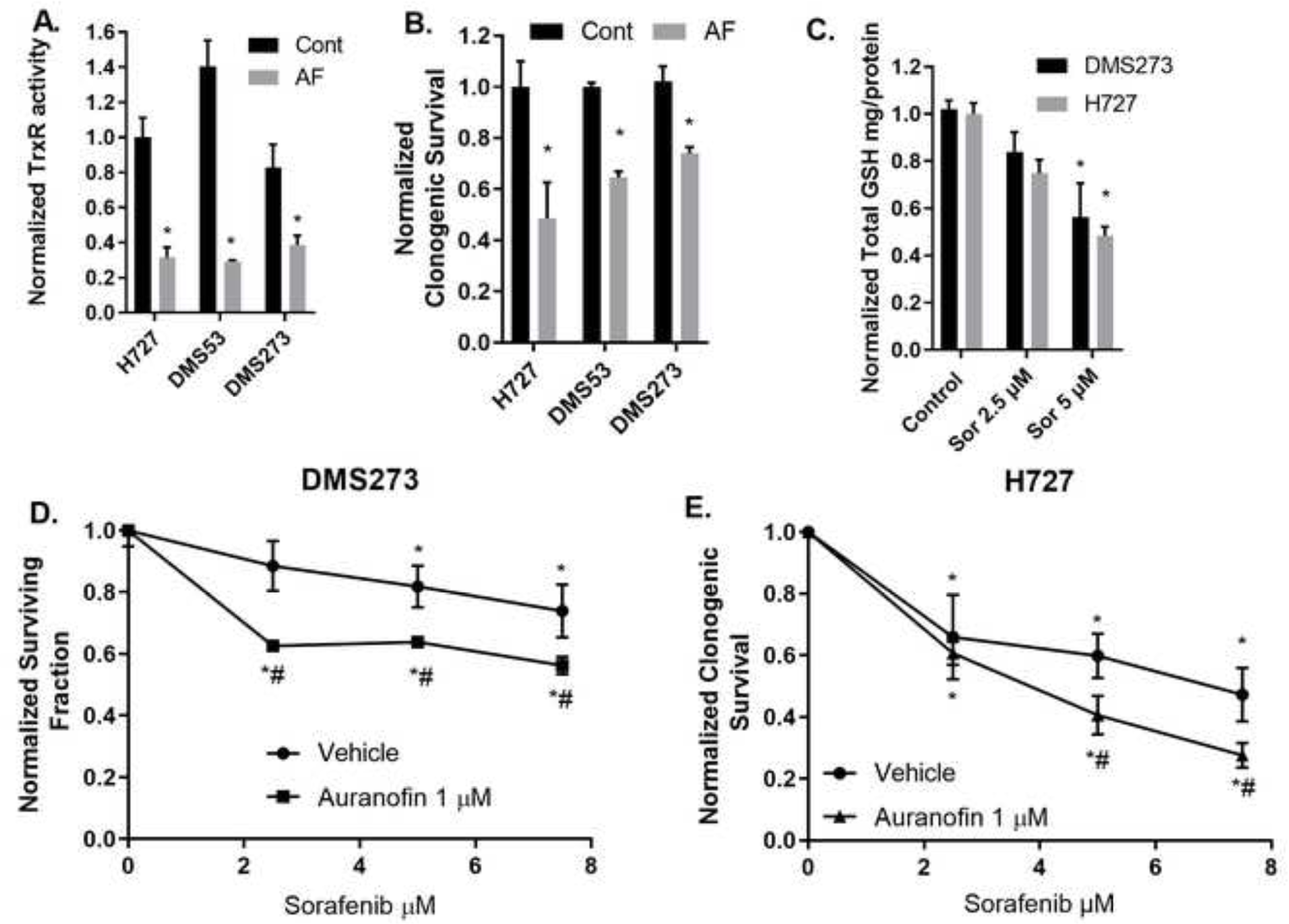
AF decreases TrxR activity and clonogenic survival of lung NET and NECs. AF enhances clonogenic cell death with sorafenib treatment. **(A)** Small cell lung cancer DMS273 and DMS53 and bronchial carcinoid H727 cell line were treated with 1 µM AF for 24 h (gray bars) then collected for TxrR enzyme activity. All activity was normalized to H727 control cells. **(B)** Same cell lines and treatment as in (A) with the clonogenic assay. Normalized to each respective control. * p<0.05 compared to untreated cells 2-way ANOVA with Fishers LSD n=3. **(C)** Exponentially growing DMS273 and H727 cells were treated for 48 h with sorafenib at 2.5 or 5 µM followed by harvest and glutathione assay. **(D,E)** DMS273 and H727 cells were treated with 48 h sorafenib and the last 24 h with 1 µM AF followed by clonogenic assay. Clonogenic survival colony counts were normalized to killing with treatment of AF alone. n ≥ 3 independent experiments 2-way ANOVA Fishers LSD * significantly different than control (p<0.05), # significantly different than either drug alone (p<0.05).

### Dosing AF to a Maximal Effective Concentration in mice

Although AF has been used historically, little is known about its pharmacokinetic and pharmacodynamic capacity in preclinical mouse models of SCLC. We sought to determine a safe and effective dose of AF in mice using a DMS273 SCLC xenograft mouse model. AF (2 or 4 mg/kg) was administered IP once (QD) or twice (BID) daily, three nude mice per group. Mice were euthanized at 24 h and 5 days after the start of injections and the TrxR activity was measured in the xenograft tumors, kidneys, and livers. 24 hours of AF administration resulted in no significant effects on TrxR activity in the tumors, but by 5 consecutive days of dosing TrxR activity was inhibited by 50% at 2 mg/kg BID, 4 mg/kg QD, and 4 mg/kg BID (**Figure 2A**). To investigate the effects of AF treatment on organs that are frequently sights of damage from cancer therapies, kidneys and livers were harvested and TrxR activity was determined. TrxR activity in kidneys was significantly reduced by 2 mg/kg QD and 4 mg/kg BID at 24 hours. TrxR activity in kidneys was significantly inhibited by 5 consecutive days of dosing in 2 mg/kg BID, 4 mg/kg QD, and 4 mg/kg BID (**Figure 2B**). TrxR inhibition in liver was only observed at an AF dose of 4 mg/kg BID at five days (**Figure 2C**).

**Figure 2.**
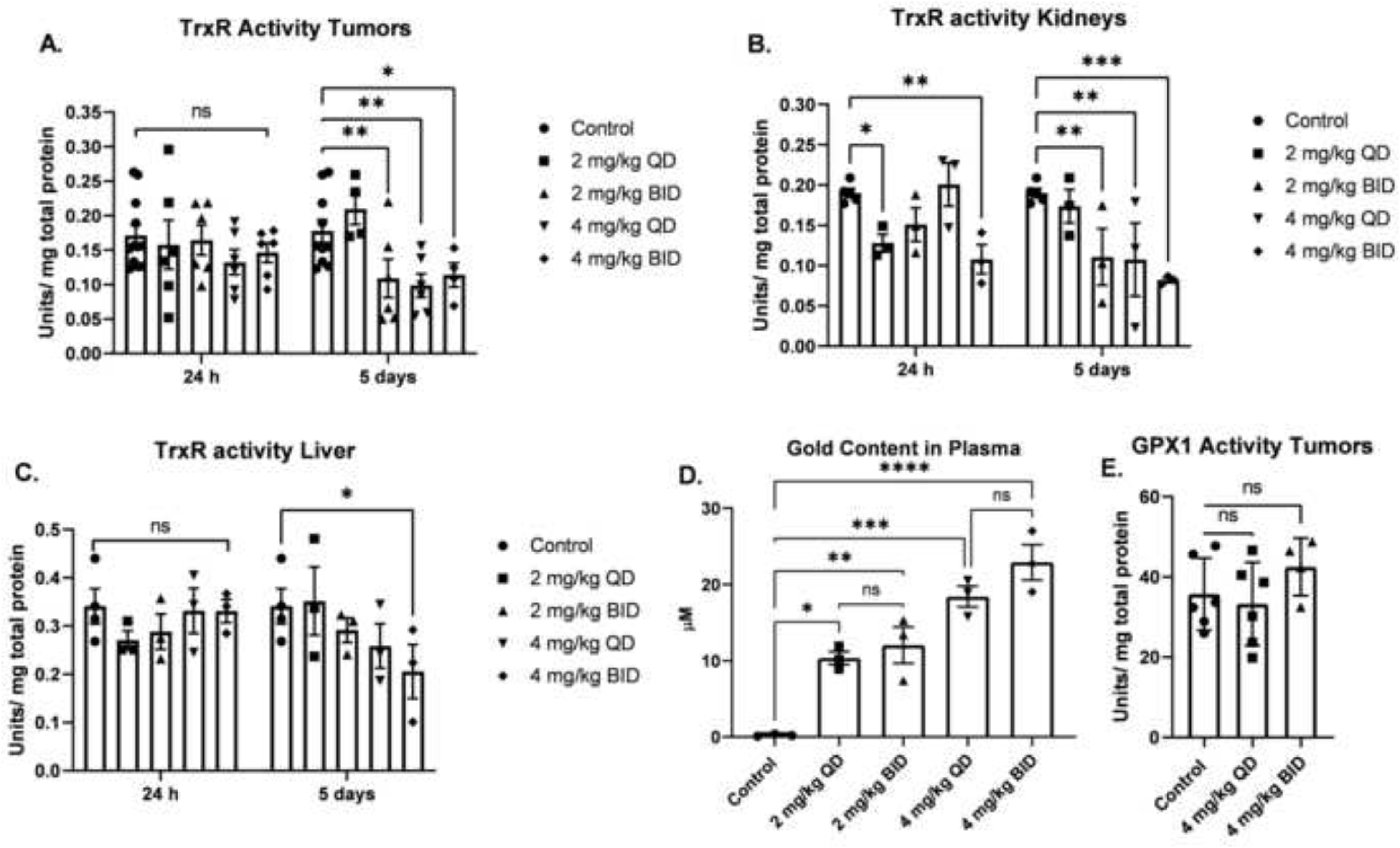
Dose escalation of AF selectively decreases TrxR activity in SCLC xenografts, kidneys and liver, but does not affect Gpx1 activity. **(A, B, C)** Female athymic nude mice (6 per group) were xenografted with 5 x10^6^ DMS273 cells (2 tumors/mouse one on each flank) and given AF IP injections for 24 h or 5 days either QD or BID. Kidney, liver, and tumor samples were collected for TrxR activity. (**D**) Plasma samples were collected 9 hours after final BID dose (5 days of therapy) and gold content determined. **(E)** Gpx1 activity was determined in xenograft tumors after 5 days of AF therapy. * = p-value < 0.05, ** = p-value < 0.01, *** = p-value < 0.001. Analyzed by Mix Effects ANOVA with Fishers LSD comparisons.

To demonstrate bioavailability of AF and to demonstrate comparisons with human blood levels, plasma gold levels were measured nine hours after the last BID dosing or 21 hours after the last QD dosing. There was a significant increase in plasma gold content of all mice treated with AF (**Figure 2D**). As expected, 4 mg/kg dosing resulted in significantly more gold in the plasma than 2 mg/kg dosing. There was not a significant difference between QD and BID dosing indicating that steady state levels of gold in the plasma can be maintained with five days of QD dosing as expected with the long half-life of AF reported in humans and rodents.^10,29^ Similar to TrxR, Gpx1 is also a selenium-containing enzyme with an active thiol moiety, and it has been reported that high doses of AF can inhibit its activity *in vitro*.^30^ GPx1 activity was measured in the tumors after five days of 4 mg/kg treatment QD and BID. Treatment with AF resulted in no significant change in GPx1 activity in the xenograft tumors (**Figure 2E**) indicating that the dose used in this study effectively inhibits TrxR activity while GPx1 is not changed *in vivo*. Together these results suggest that 4 mg/kg QD is the optimal AF dose to achieve inhibition of TrxR activity in DMS273 tumors. However, this dose also inhibited TrxR activity in the kidneys indicating the potential for nephrotoxicity when combined with other agents.

### Verifying AF inhibitory effect during typical mouse model anticancer regimens

To simulate typical anticancer dosing regimens in xenograft mouse models, the determined optimal dose of 4 mg/kg QD AF was given to mice for 14 consecutive days and then the tumors harvested. Two mice in the AF treatment group and control group (each bearing two flank tumors) were euthanized 24 hours after their final dose of AF (day 15). This regimen was repeated in the third cohort of mice with 4 mice (each with one tumor) in both the control and AF groups were again sacrificed 24 hours after the final dose of AF. 14 days of QD AF treatment significantly inhibited TrxR activity in the tumors by 75% (**Figure 3A**). Interestingly, despite the large variations of tumor size, AF was equally effective at inhibition of TrxR, confirming that AF can penetrate large tumors (**Figure 3B**).

**Figure 3.**
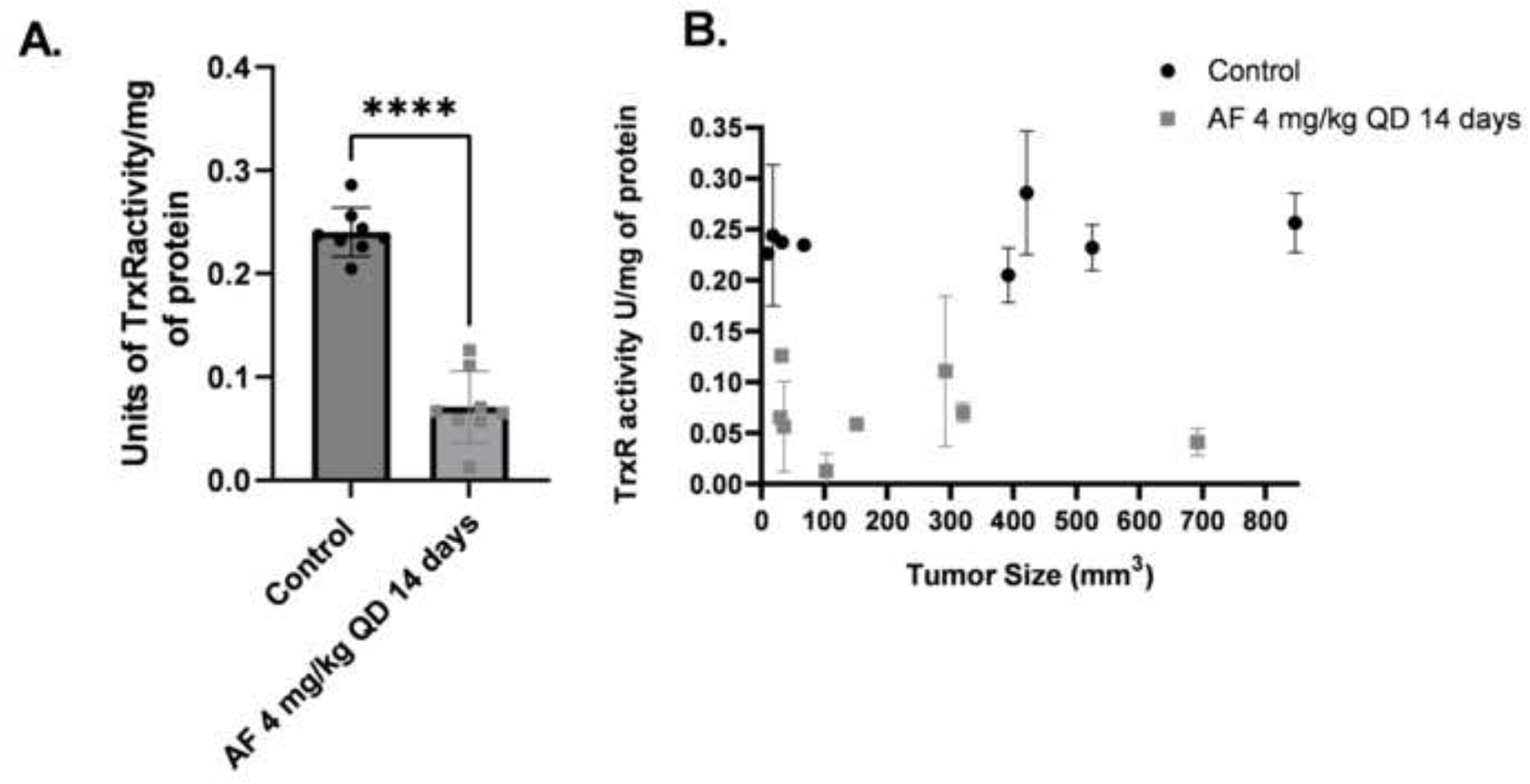
AF decreases TrxR activity in DMS273 tumors *in vivo* independent of tumor size. 2 mice each with 2 flank DMS273 xenografts in the first cohort and 4 mice with 1 flank DMS273 xenografts in the third cohort, were treated with AF 4 mg/kg IP QD for 14 days then euthanized and evaluated for TrxR enzyme activity assays and results are plotted in aggregate (A) or as a function of tumor size at the end of treatment (B). * p<0.05 compared to untreated cells two-ways ANOVA with Fishers LSD comparisons.

### AF treatment effects on tumor growth as well as animal weights, bone marrow stem cells, and blood chemistries used to assess normal tissue injury

To examine the safety and antitumor activity of AF, mice with DMS273 xenograft tumors were treated with either vehicle control or 4 mg/kg AF IP every day for 14 days. Mice were considered to have met euthanasia criteria when tumors were reached 1000 mm^3^ for 2 consecutive days This experiment was repeated on 3 separate cohorts of animals (9-10 animals/cohort) for a total of 28 AF treated mice and 30 vehicle control treated mice (supplemental Figure 1). **Figure 4A** shows % of initial weight of all the mice as a function of time after the initiation of treatment. The mice from both AF treated and vehicle control gained weight equally as well as demonstrating no signs of distress as assessed by all the general activity endpoints specified in our IACUC approved protocol. Common acute side effects of AF therapy in humans include diarrhea, skin rash and pruritus.^14^ None of these side effects were noted in any animals after the 14 days of continuous AF treatment. These results support the hypothesis that this AF dosing scheme was well-tolerated in tumor bearing mice for 14 days.

**Figure 4.**
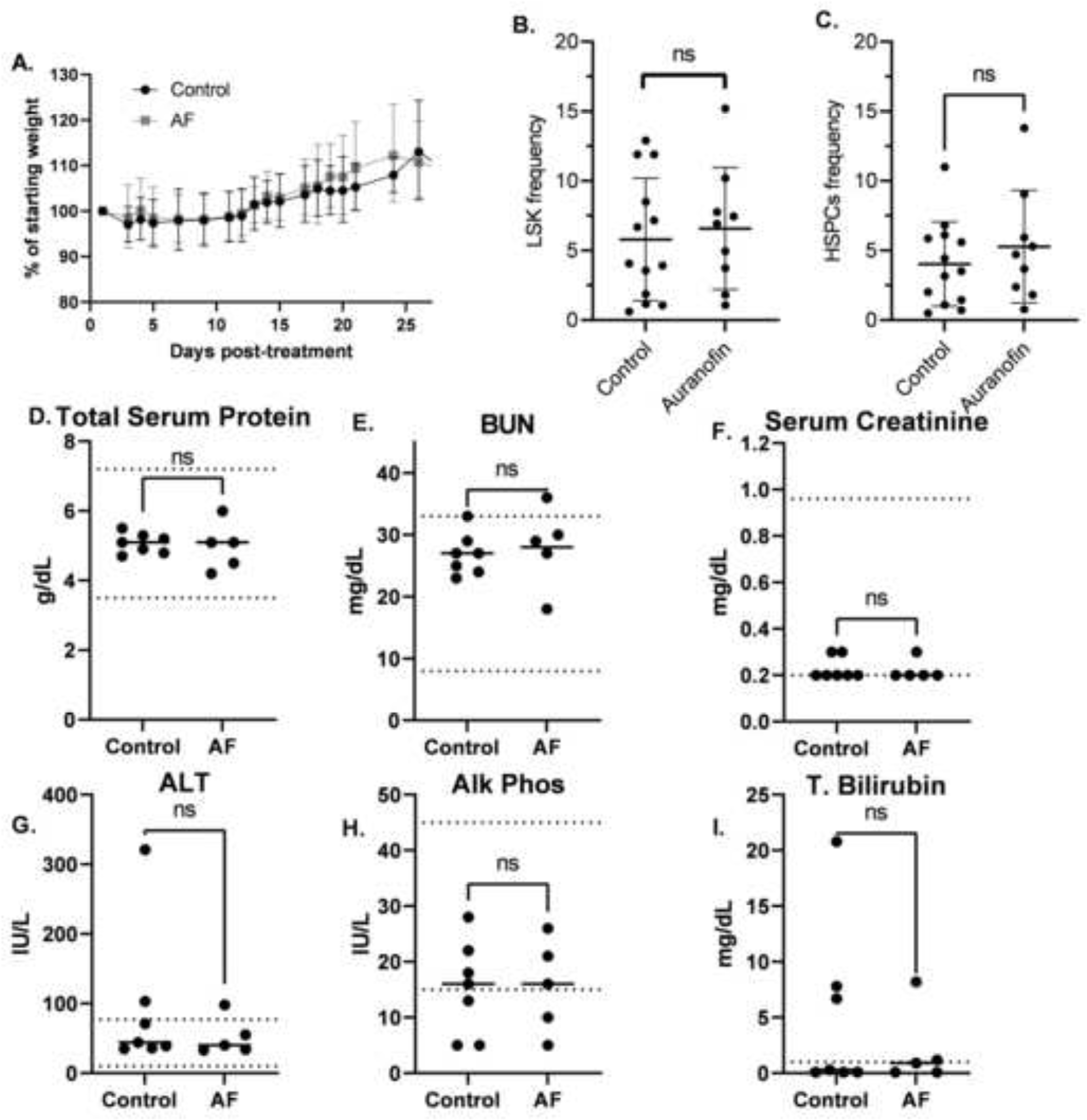
4 mg/kg QD AF IP for 14 days is non-toxic to bone marrow, liver and kidneys of mice. Female nude mice bearing DMS273 xenografts were treated with either vehicle or 4 mg/kg AF I.P every day for 14 days. Mice were euthanized when tumors were ≥ 1000 mm^3^. This experiment was done on 3 separete cohorts (9-10 animals/cohort) of animals for a total of 28 AF treated mice and 30 vehicle control treated mice. (A) Mice were monitored daily and the % of the initial weight was plotted as a function of time from the beginning of AF treatment. (**B, C**) Bone marrow from 9 AF treated mice (5 from the first cohort, 4 from the second cohort) and 13 control mice (7 from the first cohort, 6 from the second cohort) was harvested from the femurs at euthanasia and subjected to flow cytometry for HSCs (Lineage (Lin)− Sca-1+ c-Kit+, LSK) shown in (B) and HSPCs (Lin-cKit+ Sca1+ CD135-) populations shown in (C). **(D-I)** Blood was drawn from 7 control mice and 5 AF treated mice from the third cohort) via cardiac puncture at euthanasia and evaluated for clinical chemistry endpoints by Antech Diagnostics.

In humans, thrombocytopenia or leukopenia have been reported in patients on AF.^14^ When CBCs were analyzed in tumor bearing mice being treated with AF using a Siemens ADVIA 120 Hematology Analyzer (**Table 1**) there were no significant changes in WBCs, neutrophils, lymphocytes, RBCs, hemoglobin, hematocrit, mean corpuscular volume (mcv), or platelets. In order to determine any possible effects of AF on hematopoietic stem cells (HSCs and HSPCs), bone marrow from 9 AF treated mice (5 from the first cohort, 4 from the second cohort) and 13 control mice (7 from the first cohort, 6 from the second cohort) were harvested from the femurs at euthanasia and subjected to flow cytometry for HSCs (Lineage (Lin)− Sca-1+ c-Kit+, LSK) and HSPCs (Lin-cKit+ Sca1+ CD135-). **Figure 4BC** show shown no significant differences in HSCs or HSPCs with AF treatment. In addition, no significant changes of LT-HSC or ST-HSC populations were noted (data not shown). Overall, the analysis of hematological data indicated this AF dosing scheme had no adverse toxic effects on CBCs or bone marrow proliferation during the 14-day exposure.

**Table 1.** CBC analysis of peripheral blood during AF treatment. Number of mice in each group in parentheses.

|  | WBC<br>(X10<sup>3</sup>/μL) | Neutrophils<br>(X10<sup>3</sup>/μL) | Lymphocytes<br>(X10<sup>3</sup>/μL) | RBC<br>(X10<sup>6</sup>/μL) | Hemoglobin<br>(g/dl) | Hematocrit<br>(%) | MCV<br>(fl) | Platelets<br>(X10<sup>3</sup>/μL) |
| --- | --- | --- | --- | --- | --- | --- | --- | --- |
| Control (n=16) | 8.34<br>(4.39) | 2.00 (1.56) | 1.34 (1.43) | 9.15<br>(1.58) | 13.9 (2.54) | 52.6<br>(8.07) | 57.6<br>(2.06) | 1300<br>(115) |
| AF 4 mg/kg<br>(n=17) | 7.14<br>(3.50) | 1.77 (1.17) | 1.24 (1.43) | 8.82<br>(1.14) | 13.0 (1.84) | 50.5<br>(5.83) | 57.3<br>(1.60) | 1410<br>(419) |

Blood drawn from 7 control mice and 5 AF treated mice from the third cohort via cardiac puncture at euthanasia were evaluated for clinical chemistry endpoints by Antech Diagnostics (**Figure 4D-L**) and demonstrated no significant differences in total serum protein, BUN, serum creatinine, ALT, Alk Phos, or total bilirubin. These results indicated that this 14-day AF dosing scheme did not cause measurable systemic toxicity to liver or kidney.

When tumor growth (**supplemental Figure 1A-F**) and Kaplan-Meier survival curves combining all three cohorts of mice exposed to AF for 14 days were analyzed the median overall survival of the AF treated animals was found to be significantly increased (median overall survival of 19 days in control versus 23 days in AF treated animals (p=0.04) (**Figure 5**).

**Figure 5.**
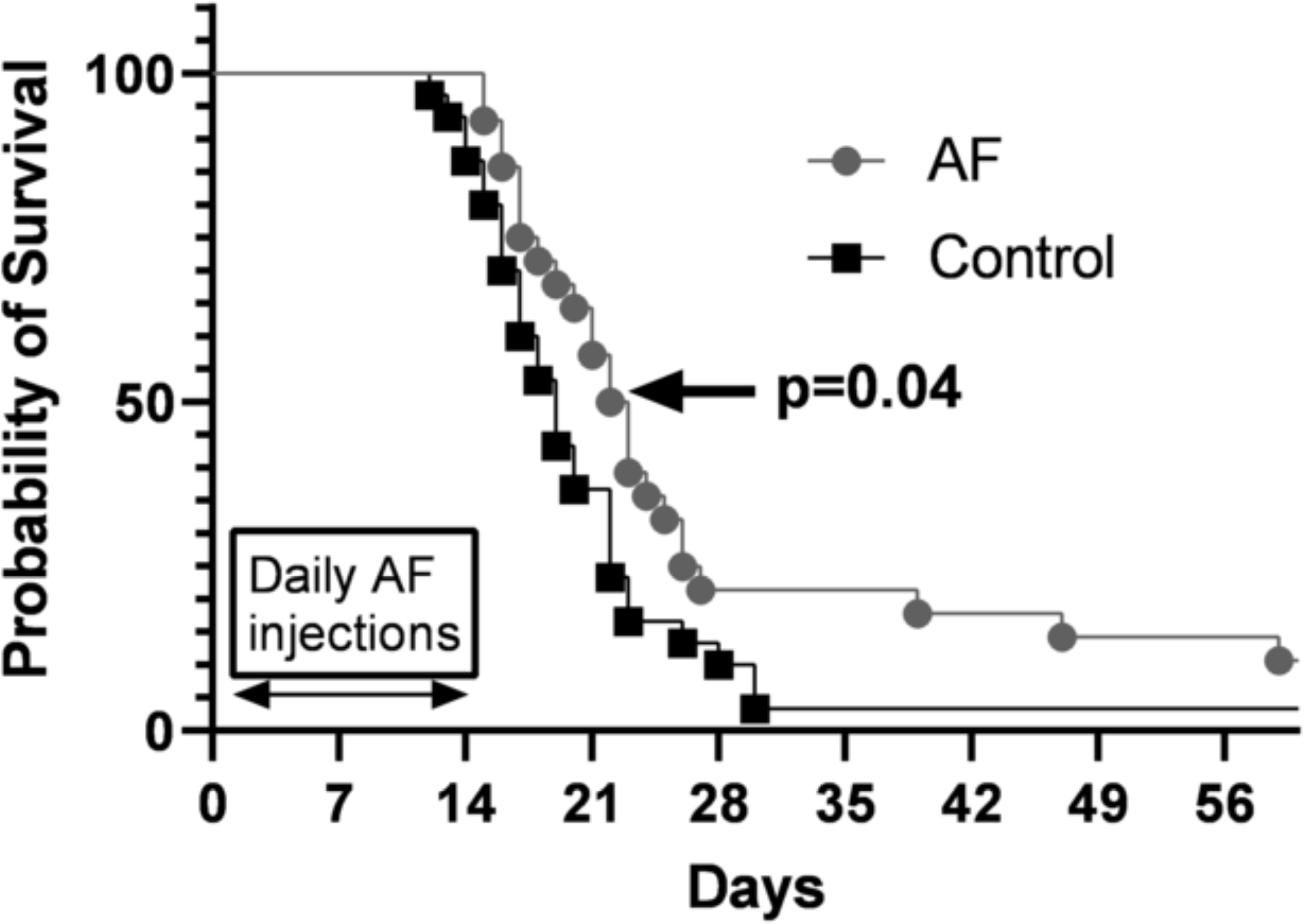
4 mg/kg QD AF IP for 14 days prolongs survival in nude mice with DMS273 xenografts. Tumors from all mice shown in **Figure 4A**, were measured via calipers and tumor volumes calculated. When tumor volumes measured 1000 mm^3^ mice were considered to have reached euthanasia criteria and Kaplan-Meier curves (three cohorts combined) were used to estimate overall survival. Log-rank (Mantel Cox) test was used to determine significance. (p<0.04)

## Discussion

AF has been FDA approved and used for rheumatoid arthritis for decades and the kinetics of the orally administered drug in humans was thoroughly studied in the 1980’s,^29,31^ by following the gold content in the patients serum. It was found that maximum gold levels were not reach for 16 weeks of treatment.^32^ An interesting question not addressed by these early studies was whether AF’s gold phosphine complex acted as a prodrug (inactive precursor) that administers the active gold molecule, or if the gold atom remains attached to the phosphine ligand during treatment. Additionally, the ability of AF to inhibit TrxR was not discovered until 1998.^16^ Targeting TrxR with AF has recently been suggested as having the potential to enhance cancer therapy responses using conventional radio-chemotherapies based on the tumor cells’ upregulation of hydroperoxide metabolic pathways that are dependent on TrxR.^5,9,17,18,26,27^ Consistent with these previous findings we have now clearly shown the AF enhances the toxicity of the broad spectrum kinase inhibitor, sorafenib, which is used as a chemotherapeutic agent in kidney cancer.^33^

Although AF has been used therapeutically for arthritis in humans for 40 years, preclinical mouse models have not been extensively studied for injury markers in the context of dosing schemes for preclinical cancer therapy studies. Consequently, we have established a once-daily, non-toxic dosing regimen of AF that effectively inhibits TrxR in a mouse SCLC xenograft model. These studies demonstrated that AF treatment resulted in dose dependent increases in plasma drug concentration that led to significant inhibition in TrxR (but not GPx1) in DMS273 xenograft tumors without any indication of weight loss, bone marrow, kidney, or liver toxicities. To compare the bioavailability of AF in mice to that of humans we measured the gold in the plasma of mice. We found the average level of the gold found in the plasma with 4 mg/kg IP dosing in mice (18.4 µM) is eleven-fold the average concentration found in humans (1.6 µM) taking the recommended dose of 6 mg/day orally for seven days for rheumatoid arthritis.^24^ The long half-life (35 days in humans) of AF suggests that higher serum and tumor drug levels can be achieved with continued dosing.^24^ In fact, the primary objective of an ongoing clinical trial (NCT01737502) is to establish the maximum tolerated dose of AF in combination with sirolimus in lung cancer, including SCLC.

In normal cells, redox metabolism is tightly coupled via non-equilibrium steady-state fluxes of reactive metabolic byproducts and electron carriers through redox sensitive signal transduction pathways. It is increasingly recognized that dysregulation of mitochondrial redox metabolism in cancer cells leads to the increased steady-state levels of reactive species including H_2_O_2_ and organic hydroperoxides that are compensated for by increased hydroperoxide metabolic pathways dependent upon NADPH and thiol containing antioxidant enzyme pathways using glutathione and thioredoxin as co-factors.^5,34^ Thioredoxin (Trx) is a small disulfide-containing protein that is maintained in a reduced state by the NADPH-dependent thioredoxin reductases (TrxR) and acts as an antioxidant both by facilitating the reduction of other proteins by cysteine thiol-disulfide exchange as well as acting as a cofactor in the peroxiredoxin-mediated detoxification of hydroperoxides.^27^ Since Trx/TrxR-dependent antioxidant pathways have been shown to be critical for cancer cells to maintain a viable steady-state in the face of increased fluxes of hydroperoxides by mitochondrial metabolism, TrxR has been proposed to be a selective metabolic target for enhancing tumor cell responses to cancer therapy.

AF was found to be an extremely potent inhibitor of mitochondrial TrxR at concentrations that are similar to those used in this manuscript. The inhibition of TrxR by AF on redox metabolism in both isolated rat heart mitochondrial redox state and intact tumor cell line redox state has been explored by Rigobello and colleagues.^15,35-38^ AF was found to be an inhibitor of isolated rat heart mitochondrial TrxR activity. In these isolated mitochondria, AF had little effect on oxygen consumption and did not alter the redox properties of the isolated mitochondrial purine nucleotides. AF treatment increased total thiol groups indicating a possible adaptive response in thiol metabolism. Also, AF (1-5 µM) increased H_2_O_2_ levels and oxidized peroxiredoxin 3 in isolated mitochondria and cells in the presence of electron transport chain inhibitors that increase ROS levels, indicating H_2_O_2_ metabolism was compromised. AF treatment increased mitochondrial total thiol groups indicating an oxidative trend.^15^ Importantly, was an increase in H_2_O_2_ in rat liver mitochondria when treated with AF in the presence of electron transport chain inhibitors. This effect depends on the decrease in the removal of H_2_O_2_ in the AF treated mitochondria.^37^ Despite the effects of AF on isolated liver and heart mitochondria noted previously, the current study show convincingly that AF had no significant toxicities in animals treated for 14 days including weight gain, liver enzymes, kidney enzymes, CBCs, or bone marrow. These results support the conclusion that during short term exposure, AF is well tolerated by normal tissues likely to be targets for conventional chemotherapies.

## Conclusion

Because tumor cells demonstrate increased levels of H_2_O_2_ and other organic hydroperoxides that is offset by increased levels of hydroperoxide metabolic pathways dependent on Trx (relative to normal cells) AF has recently been suggested as a viable adjuvant to conventional cancer therapies. Here we report the effects of AF treatment on high grade NETs and NEC and find that like several cancer types, AF decreases TrxR activity and clonogenic survival. Interestingly we also demonstrate for the first time the AF-induced enhancement of toxicity with sorafenib, a currently approved chemotherapeutic, that was also found to decrease total GSH suggesting a potential mechanism targeting both Trx and GSH mediated metabolism.

AF dosing of 14 days of 4 mg/kg daily IP in female nude mice demonstrated a highly significant 75% reduction in TrxR that was independent of tumor size indicating the AF could penetrate the tumor efficiently. In humans, common acute side effects of AF treatment include diarrhea, skin rash, proteinuria and more rarely thrombocytopenia or leukopenia. It is important to note that the AF dose used in our current study, which was effective at decreasing TrxR in SCLC xenografts, was well tolerated in mice as demonstrated by the lack of any of these common side effects of AF treatment noted in humans. There were no effects observed on the general health of the mice, weight, or changes in bone marrow progenitor cells or CBC. In addition, plasma chemistry indicated that there was no evidence of damage to the kidneys or liver. The 14-day AF dosing schedule also resulted in a modest but significant increase in median overall survival (p=0.04). Overall, the current data supports the use of AF in SCLC xenograft models as it shows little to no toxicity in mice while significantly inhibiting TrxR in the tumor. Previous work has shown that radio-chemo-sensitizers are able to improve therapy outcomes through dysregulation of oxidative metabolism ^17,26,34^, and since thoracic irradiation combined with chemotherapy continues to be a standard of care for limited stage SCLC, this work suggests the use of AF as a viable adjuvant to enhance therapeutic responses in future preclinical studies.

**Supplemental Figure 1.**
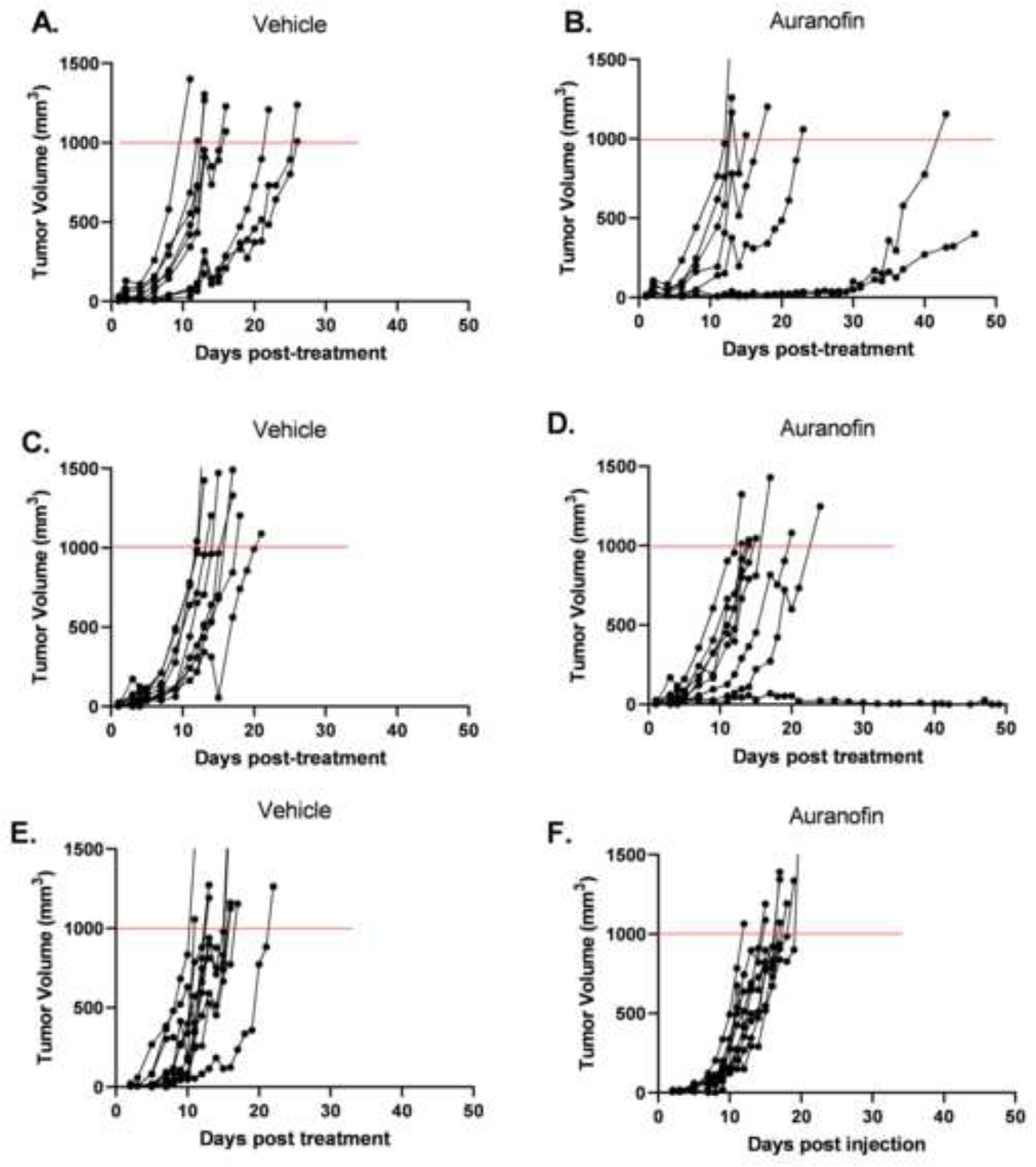
The effects of AF 4 mg/kg QD for 14 days on tumor growth. Mouse xenograft tumor volumes from the 3 cohorts of animals treated with AF and vehicle control shown in **Figure 5**, were measured via caliper and volumes plotted as a function of time (A&B for cohort 1, C&D for cohort 2, E&F for cohort 3). Mice were euthanized when their tumors measured above 1000 mm^3^. The dotted lines shows when each tumor reached 1000 mm^3^. There were no significant differences in growth rate between groups.

## Acknowledgments

We would like to thank the Holden Comprehensive Cancer Center; the University of Iowa College of Medicine Flow Cytometry Facility; the University of Iowa Office of Animal Resource; the University of Iowa Radiation and Free Radical Research Core; the Holden Comprehensive Cancer Center Biostatistical Core.

