## Supplementary figures and images for "Auranofin Inhibition of Thioredoxin Reductase Sensitizes Lung Neuroendocrine Tumor Cells (NETs) and Small Cell Lung Cancer (SCLC) Cells to Sorafenib as well as Inhibiting SCLC Xenograft Growth"

### supplimental figure 1

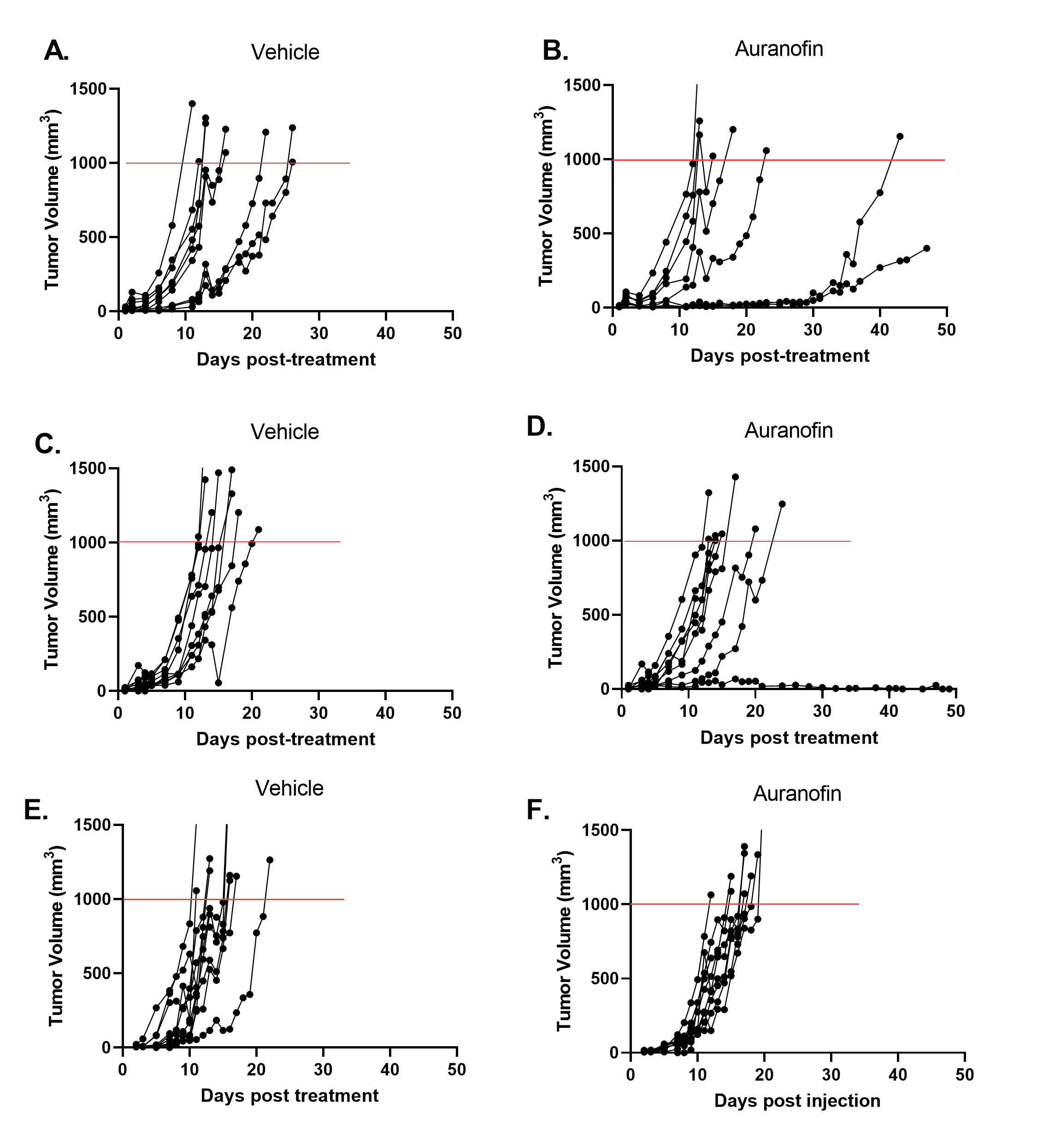
